# Intercellular Communication via the *comX*-Inducing Peptide (XIP) of *Streptococcus Mutans*

**DOI:** 10.1101/148320

**Authors:** Justin Kaspar, Simon A. M. Underhill, Robert C. Shields, Adrian Reyes, Suzanne Rosenzweig, Stephen J. Hagen, Robert A. Burne

## Abstract

Gram-positive bacteria utilize exported peptides to coordinate genetic and physiological processes required for biofilm formation, stress responses and ecological competitiveness. One example is activation of natural genetic competence by ComR and the *comX-*inducing peptide (XIP) in *Streptococcus mutans*. Although the competence pathway can be activated by addition of synthetic XIP in defined medium, the hypothesis that XIP is able to function as an intercellular signal molecule has not been rigorously tested. Co-culture model systems were developed that included a “sender” strain that overexpressed the XIP precursor (ComS) and a “responder” strain harboring a GFP reporter fusion to a ComR-activated gene (*comX*) promoter. The ability of the sender strain to provide a signal to activate GFP expression was monitored at the individual cell and population levels using i) planktonic culture systems, ii) cells suspended in an agarose matrix or iii) cells growing in biofilms. XIP was shown to be freely diffusible and XIP signaling between the *S. mutans* sender and responder strains did not require cell-to-cell contact. The presence of a sucrose-derived exopolysaccharide matrix diminished the efficiency of XIP signaling in biofilms, possibly by affecting spatial distribution of XIP senders and potential responders. Intercellular signaling was greatly impaired in a strain lacking the primary autolysin, AtlA, and was substantially greater when the sender strain underwent lysis. Collectively, these data provide evidence that *S. mutans* XIP can indeed function as a peptide signal between cells and highlight the importance of studying signaling with endogenously-produced peptide(s) in populations in various environments and physiologic states.

**IMPORTANCE:** The *comX*-inducing peptide (XIP) of *Streptococcus mutans* is a key regulatory element in the activation of genetic competence, which allows cells to take up extracellular DNA. XIP has been found in cell culture fluids and addition of synthetic XIP to physiologically receptive cells can robustly induce competence gene expression. However, there is a lack of consensus as to whether XIP can function as an intercellular communication signal. Here, we show that XIP indeed signals between cells in *S. mutans*, but that cell lysis may be a critical factor, as opposed to a dedicated secretion/processing system, in allowing for release of XIP into the environment. The results have important implications in the context of the ecology, virulence and evolution of a ubiquitous human pathogen and related organisms.

## INTRODUCTION

Evidence for extensive complexity and diversity in bacterial intercellular communication strategies has rapidly accumulated since quorum sensing was first described by Greenberg and colleagues (1). The study of the genetics, biochemistry and ecology of interbacterial communication has yielded valuable insights into the molecular basis for the behaviors of bacterial communities and has revealed novel pathways that can be targeted to disrupt the formation, persistence and pathogenic potential of biofilms (2). Because bacterial biofilm communities exist at relatively high cell densities and are typically rich in exopolymeric materials, mass transport limitations contribute to the development of considerable spatial heterogeneity. Despite this heterogeneity, biofilm communities coordinate population-wide responses when challenged with exogenous and endogenous stressors by employing intercellular communication pathways. In the well-characterized Gram-negative quorum sensing systems, cell-cell communication occurs primarily via N-acyl homoserine lactones (3), whereas Gram-positive bacteria communicate mainly via oligopeptides, often through two-component signal transduction systems (TCS) consisting minimally of a membrane-associated histidine kinase and a cytosolic response regulator (4). Alternatively, signaling may occur through a pathway that requires active internalization of a peptide, followed by specific binding of the peptide by a cytosolic transcriptional regulator (5). Genetic competence in Gram-positive bacteria is usually controlled by one of two peptide-based systems. The most thoroughly studied are similar to the ComCDE pathway of *Streptococcus pneumoniae.* ComC is the pro-peptide of competence stimulating peptide (CSP), which is processed and exported by a specialized secretion apparatus, then signals for the activation of competence genes via the ComDE TCS. The second is the more recently characterized ComRS pathway, which consists of the ComS precursor for *comX-*inducing peptide (XIP) and the cytosolic regulator ComR (6).

*Streptococcus mutans*, which colonizes the human oral cavity, is unusual among streptococci in that both CSP and XIP can activate transcription of the master regulator of competence development, an alternative sigma factor encoded by the *sigX* (σ^X^) or *comX* (7–9) gene; SigX and ComX are homologues in different organisms. Thus, *S. mutans* has become an intriguing model for the study of peptide-based communication strategies (10). CSP in *S. mutans* is generated by processing and secretion of ComC through the ComAB exporter, with further processing by the SepM protease to yield the most active form (18-aa) of CSP (11). In a chemically-complex medium that is rich in peptides, such as brain heart infusion (BHI), CSP signaling occurs via ComDE and directly activates bacteriocin production. However, ComCDE of *S. mutans* are not true homologues of *S. pneumoniae* ComCDE, rather they are most similar in sequence and function to the BlpCRH system of the pneumococcus, with which *S. mutans* ComCDE share ancestral origin (6, 9). Provision of CSP to early exponential phase cells results in up-regulation of *comX* transcription through an as-yet-undefined indirect mechanism, and leads to a dramatic increase (~10^3^-fold) in efficiency of transformation. The second route for induction of competence in *S. mutans* involves the ComRS pathway, consisting of the Rgg-like transcriptional regulator ComR and XIP, the latter being a 7-aa peptide derived from the C-terminus of the 17-aa precursor, ComS. The current view of ComRS-dependent activation of *comX* in *S. mutans* is that the ribosomally translated ComS peptide is exported into the extracellular space and cleaved by a protease(s) to yield XIP (7, 9, 12), although the mechanisms for secretion or processing have not been identified. Notably, three independent research groups have detected XIP in supernatant fluids of *S. mutans* (13–15). In *Streptococcus thermophilus* (16), the Eep protease processes ComS to XIP, but an equivalent function for proteases in *S. mutans* with characteristics similar to Eep has not been demonstrated (13). Additionally, in *S. thermophilus*, the exported XIP appears to remain associated with the cell surface, as intercellular signaling by XIP in a co-culture system required cell-cell contact (16). The current model for how the ComRS-XIP system functions is that, after XIP is produced and released, it is transported back into the cell by the oligopeptide ABC transporter Opp (9). The re-imported XIP can then be specifically bound by ComR to form a dimeric ComR-XIP complex that functions as a transcriptional activator of the promoters of *comX* and *comS* (17). The activation of *comS* creates a positive feedback loop that amplifies ComS and possibly XIP production (9, 17). The ComR-XIP complex recognizes a ComR-box consisting of a 20-bp palindromic motif with a conserved central GACA/TGTC inverted repeat (7, 18). The degree of conservation of the ComR-box with the consensus sequence has been correlated with levels of ComR-regulon expression (17).

Several different orthologs of ComR and ComS are present within streptococci (19). ComR, along with apparent orthologous proteins that include PlcR of *Bacillus thuringiensis* and PrgX of *Enterococcus faecalis*, are part of the RNPP superfamily (Rap/Npr/PlcR/PrgX) of transcriptional regulators that interact with pheromones in cell-cell signaling pathways. Rgg regulators typically contain an N-terminal helix-turn-helix (HTH) DNA-binding element and an approximately 220-aa C-terminal alpha-helical domain thought to be involved in binding of cognate small hydrophobic peptides (SHPs) (20). ComS peptides show substantial size and primary sequence variation between species of streptococci (21, 22). The so-called type I ComS peptides of *Streptococcus salivarius* and *S. thermophilus* harbor a P(F/Y)F motif and lack charged residues. The type II peptides, like ComS of *S. mutans*, along with the Pyogenes and Bovis groups of streptococci, contain a WW motif and basic and/or acidic residues (7, 17). In all streptococci that possess ComRS, *comS* is located immediately downstream of *comR*. The ComR-box upstream of *comS* is immediately followed by a “T-tract” that, in conjunction with the ComR-box palindrome, may function as a Rho-independent transcriptional terminator for *comR* (7, 17). All type II ComRS systems, including that of *S. mutans*, are located about 50 kbp from the origin of replication and are associated with clusters of genes involved in purine biosynthesis and the RuvB Holliday junction DNA helicase (7).

In the period since the ComRS system was first described by Gardan (23), Mashburn-Warren (7) and their co-workers, there has been minimal investigation into how XIP is released and whether it can actually function as an intercellular communication molecule for *S. mutans* and related organisms. In this report, we designed and constructed an *S. mutans* co-culture system that allowed us to investigate intercellular ComRS signaling through observation of individual cells in planktonic cultures, in an agarose gel matrix and within single-species biofilm communities.

## RESULTS AND DISCUSSION

### Development of in vitro co-cultivation models to monitor XIP signaling

While numerous studies have used synthetic XIP peptide (sXIP) to explore the activation of *the* ComRS signaling pathway of *S. mutans* in planktonic cultures, little attention has been devoted to self-activation of the system, and only recently has sXIP-dependent gene activation been examined in *S. mutans* growing in biofilms (24), which is the normal growth mode of this organism in the oral cavity. Here, we developed a system to test whether the ComRS system can self-activate (intracellular) or cross-activate (intercellular) *comX* in the absence of exogenously supplied sXIP. Using previously described strains for our studies of genetic competence in *S. mutans* (25–27), we constructed a co-cultivation model consisting of two genetically modified *S. mutans*: a “sender” strain harboring a *comS* overexpressing plasmid (pIB184comS) along with a plasmid carrying the dsRed fluorescent protein (RFP) under the control of the *comX* promoter (P*comX*), and a “responder” strain expressing GFP also under the control of P*comX* (Figure 1A). The responder also carried the empty pIB184 vector so that the two strains were as genetically similar as possible. In certain cases, *comS* was deleted from the responder to remove any confounding effects of XIP production and auto-feedback by the responder strain. As structured, then, the sender constitutively expresses *comS* from the P_23_ promoter on pIB184, overproducing the pro-peptide ComS and hence XIP. Intracellular signaling or self-activation could be monitored in the sender strain through RFP fluorescence; the sender strain was easily distinguished from the co-cultivated responder in microscopy images by its lack of green fluorescence in combination with its strong red fluorescence. If XIP can serve as an intercellular signal, the responder strain should then import the XIP from the sender strain using Opp, where it can complex with the ComR regulator and activate the P*comX* promoter, resulting in production of GFP. Intercellular signaling can be visualized by fluorescence microscopy, by measuring total fluorescence in a plate reader or by flow cytometry of both planktonic cultures and/or dispersed biofilms.

**Figure 1.**
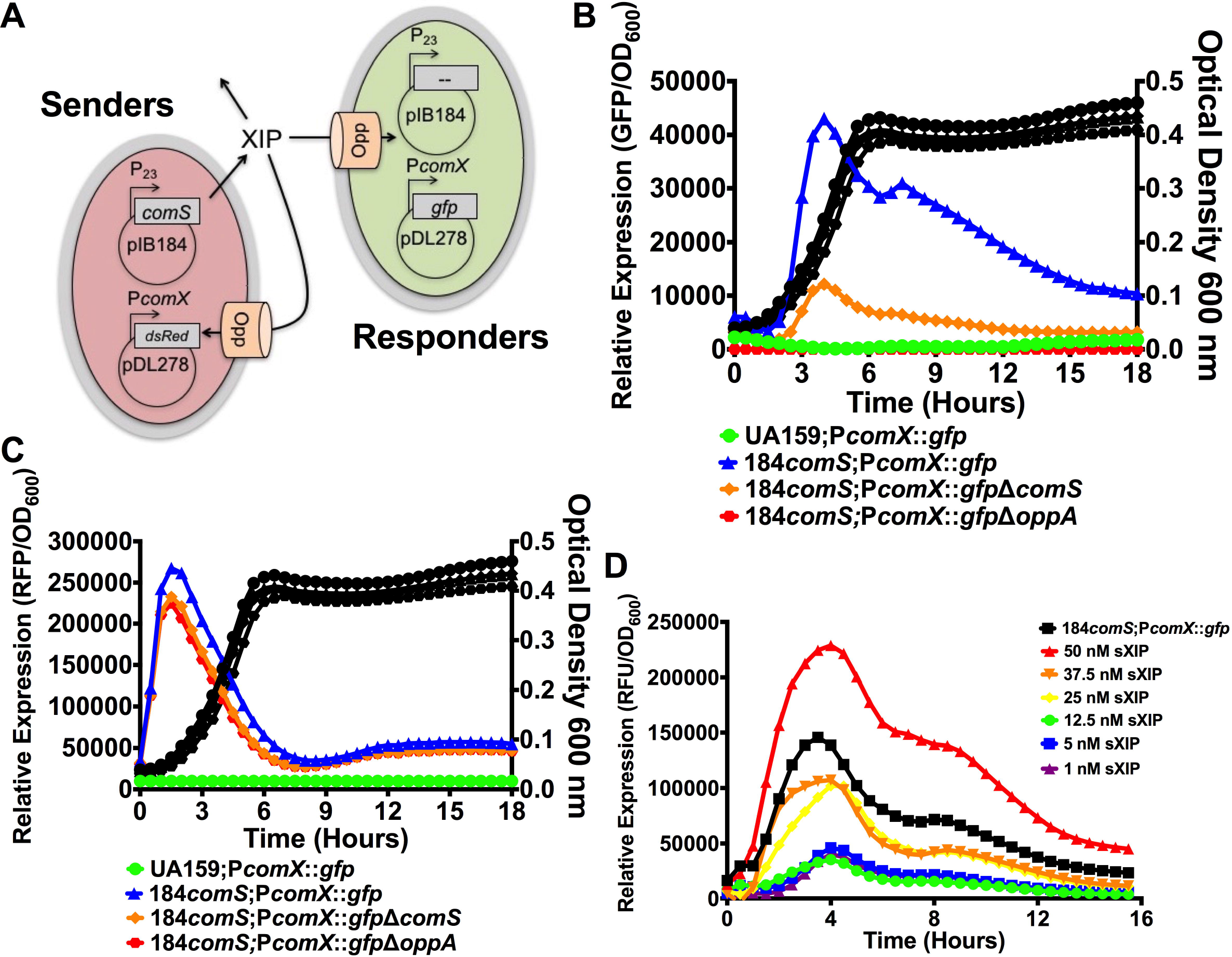
Model of co-culture system. (A) To monitor natural ComRS signaling, two strains of *S. mutans* are grown together in a co-culture system. The first strain, the “sender”, contains a plasmid that allows for the overexpression of the XIP peptide precursor *comS* under the control of the constitutive P_23_ promoter. The sender strain also contains pDL278 carrying the gene for the dsRed fluorescent protein under the control of the *comX* promoter. The second strain, the “responder”, harbors the P*comX*::*gfp* reporter plasmid on pDL278 which becomes activated when external XIP is imported into the responder via the oligopeptide permease, Opp. The empty pIB184 vector is also harbored in the responder strain to keep sender and responder strains as genetically similar as possible. (B) Relative GFP expression and (C) relative RFP expression with OD_600_ measurements during co-culture growth of UA159 and P*comX*::*gfp*/UA159 (green; circles), pIB184comS/UA159 and P*comX*::*gfp*/UA159 (blue; triangles), pIB184comS/UA159 and P*comX*::*gfp*/ΔcomS(orange,diamonds)and P*comX*::*gfp*/Δ*oppA* (red; squares). (D) Relative GFP expression measurements during co-culture of UA159 (sender) and P*comX*::*gfp*/UA159 (responder) using different concentrations of sXIP (legend) in comparison to growth between pIB184comS/UA159 and P*comX*::*gfp*/UA159 (black squares; co-culture). Each assay was performed with biological triplicates.

The ability of the sender strain to activate *comX* in the responder strain, as measured by GFP production, was first examined with a plate reader-based assay. Mid-exponential phase planktonic cultures of both sender and responder strains were diluted 1:100 into fresh FMC medium, such that the sender strain was present in a ratio of 2.3:1 with the responder strain (Supplemental Figure 1), resulting in a final overall dilution in terms of absolute cell numbers of approximately 1:50. After 18 hours of growth, no detectable fluorescence was observed when wild-type (WT) *S. mutans* strain UA159 was co-cultivated with the P*comX*::*gfp* responder strain (Figure 1B). In contrast, the *comS*-overexpressing strain, i.e. the sender, was able to elicit high expression of P*comX*::*gfp* by the responder under similar growth conditions. To verify that P*comX*::*gfp* expression was derived from imported extracellular XIP provided by the sender strain, we evaluated GFP fluorescence with two mutant backgrounds of the responder strain, Δ*comS* and Δ*opp*. GFP fluorescence was still detectable in the responder strain lacking *comS*, albeit at a level 4-fold lower than in the strain with an intact *comS* gene. However, no GFP production was observed in the *opp* mutant of the responder strain. Both results were as expected: a lower level of fluorescence in a responder lacking *comS* would be unable to amplify *comX* expression through the internal positive feedback arising from the ComR-XIP complex activating the *comS* promoter, and XIP provided by the sender must be imported via Opp to activate *comX*. In terms of the sender strain, P*comS* feedback via the RFP reporter was measured with the same intensity and timescale between all ComS-over production strains, suggesting that the observable responder differences were due to their genetic background alone (Figure 1C). Collectively, the results indicated that the sender was able provide a XIP signal to the responder and that *comX* was activated by this XIP signal, consistent with the current model for XIP-dependent activation of *comX* via active internalization by Opp, complexation with ComR, and signal amplification through activation of *comS* transcription.

To provide a rough estimate of the amount of XIP generated by the sender strain, we compared the fluorescence of the pIB184ComS/UA159;P*comX*::*gfp*/UA159 co-cultivation results to the fluorescence of cultures obtained by addition of various concentrations of sXIP to a mixed culture of UA159 and P*comX*::*gfp*/UA159 at the same 2.3:1 ratio used in the previous experiment. From the measured fluorescence, it could be estimated that the amount of XIP provided by the *comS* over-expresser was between 37.5 and 50 nM (Figure 1D), which is similar to the concentration of XIP required to achieve ComRS-dependent activation of *comX* in other *S. mutans* UA159 derivatives in microfluidic and planktonic experiments (9). These data collectively show not only that intercellular signaling can occur via XIP and the ComRS pathway in *S. mutans*, but also that signaling can occur in the co-culture model at physiologically relevant levels of the peptide signal.

### XIP signaling does not require cell-cell contact

Early studies concluded that *S. mutans* XIP was a secreted, diffusible signal molecule (14). However, studies with *S. thermophilus* demonstrated that cell-cell contact was required for intercellular signaling by XIP (16). Since *S. mutans* and *S. thermophilus* are not closely related within the genus *Streptococcus* and evolved in very different environments, we tested whether the signaling observed in Figure 1 required cell-cell contact. Our initial test involved determining whether exposure of the responder strain to cell-free supernatant fluids derived from the sender strain could induce *comX* expression. Overnight cultures of either UA159 or the *comS*-overexpressing strain were grown in FMC medium. After pelleting the cells, the supernates were filter sterilized, the pH was adjusted to 7.0, and a concentrated solution of sterile glucose was added to provide an additional 20 mM glucose. Supernates from *S. mutans* UA159 supplemented with 50 nM sXIP served as a positive control. The P*comX*::*gfp*/UA159 responder was then suspended in the supernatant fluids and fluorescence was measured during growth of the cells. No fluorescence was observed from the responder strain grown in the supernates from UA159, unless sXIP was added (Figure 2A). However, when the P*comX*::*gfp* responder strain was grown in supernates from the *comS*-overexpressing strain, robust fluorescence was evident. These data clearly show that cell-free supernates of *S. mutans* are sufficient for intercellular signaling, albeit overexpression of *comS* was required to yield sufficient signal peptide in the supernates under the conditions tested.

**Figure 2.**
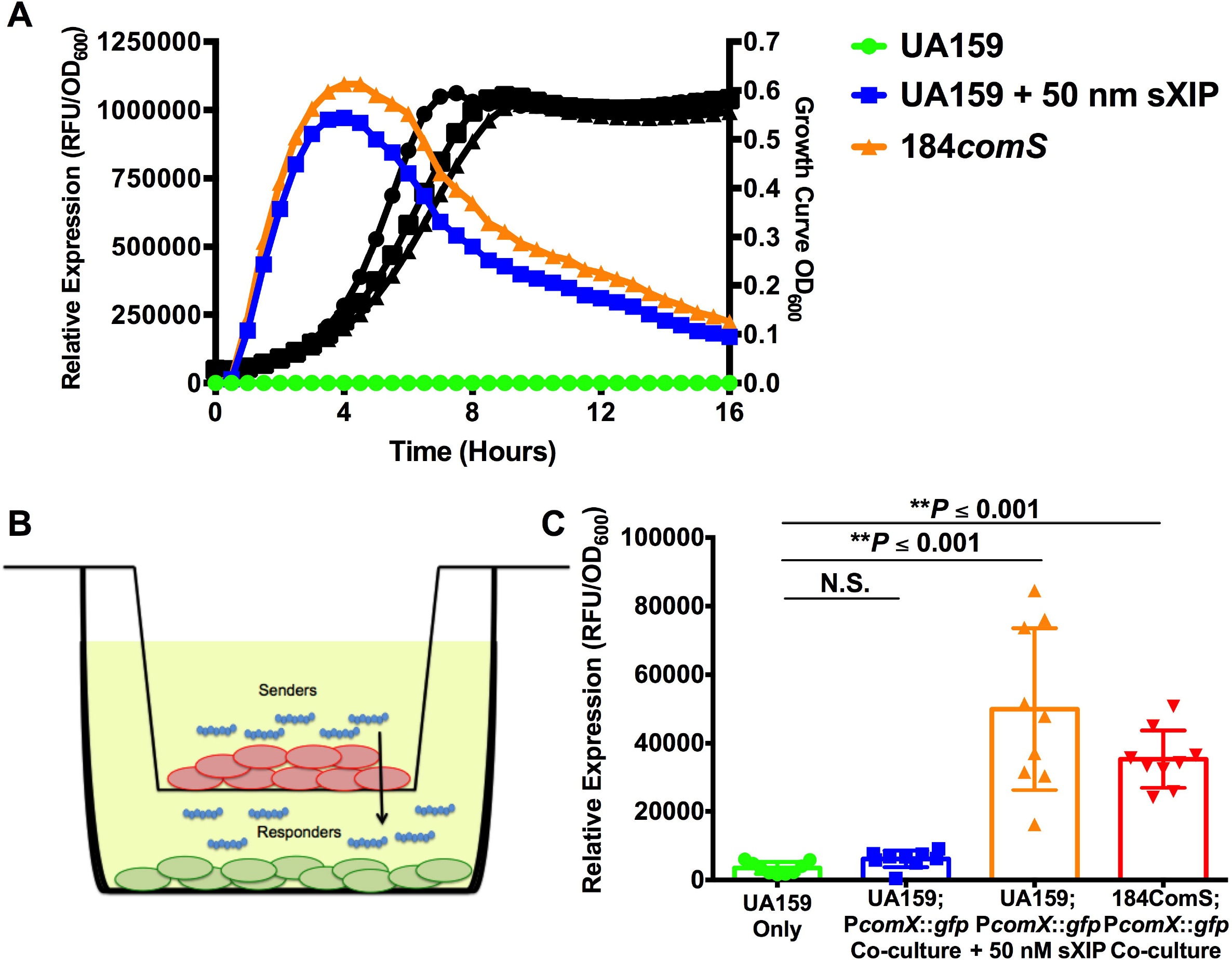
XIP signaling is cell-cell contact independent. (A) Relative GFP expression and OD_600_ measurements of P*comX*::*gfp*/UA159 grown either in supernatants of either UA159 (green, circles), UA159 supplemented with 50 nM sXIP (blue, squares), or pIB184comS/UA159 (orange, triangles). Supernatants were taken from overnight cultures of respective strains and filter sterilized, its pH was adjusted to 7.0, and glucose was added to the spent medium in an amount equivalent to a concentration of 20 mM. (B) Model of transwell assay using co-cultures of pIB184comS/UA159 and P*comX*::*gfp*/UA159. P*comX*::*gfp*/UA159 (green) strain is inoculated first in the bottom well, followed by placem ent of the 0.4 μM polycarbonate membrane insert The pIB184comS/UA159 (red) strain is then inoculated in the top well above the membrane. (C) Relative GFP expression measurements from transwell experiments. Labeling of x-axis denotes strain inoculated in the upper well first, followed up the strain inoculated in the lower well. Each assay was performed with biological triplicates. Statistical analysis was performed by Student’s *t*-test. N.S. = not significant.

Definitive evidence that the sender could produce a cell-free signal that could activate *comX* expression in the responder strain was obtained when the *comS*-overexpressing sender and PcomX::*gfp*/UA159 responder strains were cultured in separate chambers of a transwell apparatus (Figure 2B), with the strains separated by a 10 µm thick, 0.4 μm pore-size polycarbonate filter membrane. The membrane allows passage of small molecules and peptides, but prevents physical contact between cells in the different compartments. Addition of 50 nM of sXIP to medium alone in the upper compartment activated P*comX*::*gfp* expression in the lower compartment (Figure 2C). Consistent with the supernatant transfer experiments in Figure 2A, UA159 was unable to stimulate *comX* expression in the P*comX*::*gfp* responder over the course of a 24-h incubation. In contrast, the *comS*-overexpressing strain in the top compartment of the transwell apparatus readily stimulated *comX* expression in the responder strain in the lower compartment. These experiments show that signaling occurs through the polycarbonate filter membrane and does not absolutely require contact, which suggests that XIP is released by the sender strain, is diffusible and that signaling can occur without direct contact between sender and responder strains of *S. mutans*.

In previous reports, the XIP peptide could be detected in supernatant fractions of the wild-type strain UA159 at a high cell densit*y* (OD_600_ = 1.0) and cell supernates could also stimulate a P*comX* reporter strain (14). In this study, we were not able to observe activation of the P*comX*::*gfp* reporter by co-cultivation of strain UA159 or by using supernates from overnight cultures of *S. mutans* UA159. The discrepancy between these studies and ours could be due to the previously documented differences in XIP signaling seen between the chemically-defined media CDM (14) and FMC (28). In particular, XIP signaling is exquisitely sensitive to low pH (28, 29) and the drop of pH in FMC due to carbohydrate metabolism may be too rapid when compared to that in CDM, since the latter is formulated with substantially greater (phosphate) buffer capacity. While levels of XIP in supernates from high-density, overnight cultures have been reported to be as high 1 μM (14), such levels are not required to activate *comX* transcription or induce genetic competence in early exponential phase cells when the pH is near neutrality (28, 29) (Figure 1C). In fact, later in the growth phase, when XIP concentrations in the medium may approach μM levels, the ComRS system may be inactive due to acidification of the environment. Instead, optimal signaling occurs at the threshold for P*comX* activation and theoretically at lower cell densities, when the inhibitory effects of low pH generated by carbohydrate fermentation are minimalized. Thus, the co-culture model that is presented here permits the study of competence signaling under conditions that may be more physiologically relevant.

### XIP diffuses in an agarose matrix

Oral biofilms, the natural habitat of *S. mutans*, are rich in exopolymeric material of bacterial and host origin. To verify that the XIP is able to diffuse freely in an aqueous environment containing *S. mutans*, we conducted a diffusion experiment using the P*comX*::*gfp*/Δ*comS* reporter strain grown in FMC medium and embedded in low-melting-point agarose/FMC medium. Cells were loaded into an IBIDI microslide channel (IBIDI GmbH) and allowed to attach to the channel surface. A 2% low-melting point agarose/FMC mixture was then injected into the channel to prevent advective transport of XIP and to immobilize the cells in the channel. sXIP, diluted in FMC to a final concentration of 1 μM, was then deposited at one end of the channel with an equal volume of water deposited at the other end to balance the hydrostatic pressure along the length of the channel. Cells at different locations in the channel were then imaged by fluorescence microscopy at regular time intervals as the XIP diffused through the channel (Figure 3B-D). The diffusion coefficient of XIP was estimated by modeling the channel as one-dimensional with a concentrated XIP source at one end from which the peptide spreads diffusively (that is, the cells produce no XIP and there is no hydrodynamic flow). GFP fluorescence versus time and spatial position was fit to a 1D diffusion equation (Figure 3A), leading to an estimate for the diffusion coefficient of XIP through FMC/agarose. The resulting estimate of 1.8 ± 0.3 x 10^−6^ cm^2^/s was of the expected magnitude, based on the molecular mass of XIP and diffusion in an aqueous medium. These data, along with the transwell experiment presented in Figure 2, verify that XIP can signal by diffusion through aqueous medium, independently of cell-cell contact.

**Figure 3.**
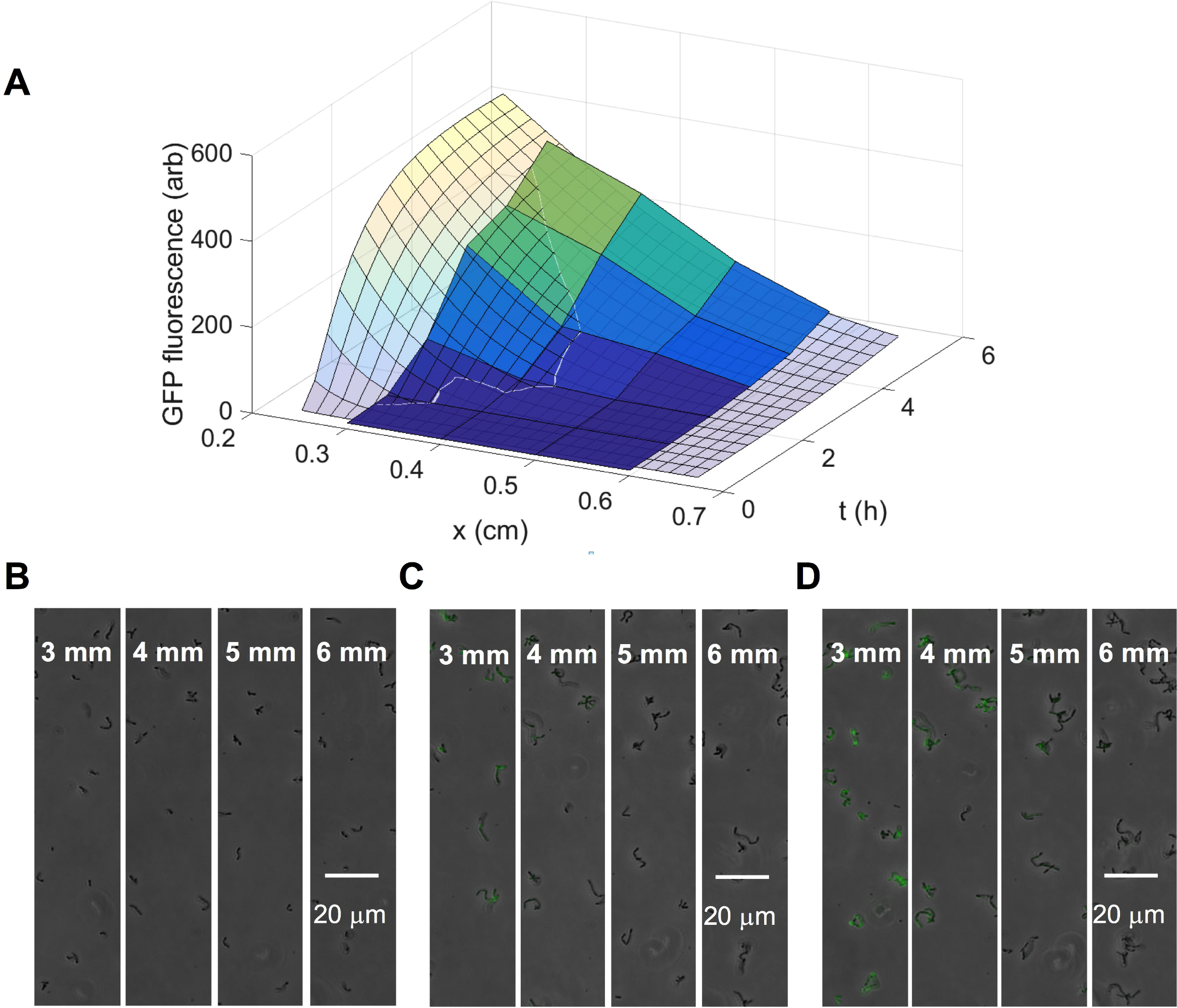
Measurements of XIP spatial diffusion. P*comX*::*gfp*/Δ *comS* cells were injected into an IBIDI microslide and allowed to settle before an agarose/FMC mixture was pushed through the slide to immobilize the cells. sXIP (1 μM) was injected into one end of the channel and green fluorescence of individual cells was monitored at different distances from the site of XIP injection. (A) 3D plot of average GFP fluorescence of cells in the channel (solid) overlaid with a fit to a diffusive spreading model (mesh). (B) Phase/fluorescence overlaid images of P*comX*::*gfp* Δ *comS* at distances of 3, 4, 5 and 6 mm from the XIP injection site at 1 h after injection; (C) 3 h after injection; (D) 5 h after injection.

### Study of ComRS signaling between individual cells in an agarose matrix

To begin visualizing ComRS signaling at the single-cell level using a co-culture approach, we first mixed the *comS*-overexpressing sender with the responder, which carried P*comX*::*gfp* in either the UA159 or Δ*comS* genetic background, in liquid FMC when the cells reached an OD_600_ = 0.1. The mixture was loaded into a microfluidic channel, followed by injection of the 2% low-melting-point agarose/FMC mixture described above and in Figure 3. Both the sender and the responder were imaged at 1-h intervals for up to 5 h to monitor intercellular activation of P*comX* via diffusion of XIP. Figure 4 plots the measured fluorescence of both RFP and GFP at the 2 h and 3 h time points. The scatter plots of individual cell fluorescence in the culture are shown for both a UA159-based responder (Figures 4A and 4E) and for a Δ *comS* responder (Figures 4C and 4G). In both co-cultures, P*comX*::*gfp* activation in the responders was not significantly different from baseline fluorescence of the wild-type strain UA159, which lacks a *gfp* reporter (data not shown). The experiment was repeated with a positive control in which 1 μM sXIP was added into co-cultures containing responders with a UA159 (Figures 4B and 4F) or Δ*comS* genetic background (Figures 4D and 4H). In this case, most cells produced either red fluorescence (senders) or green fluorescence (responders), but not both. This was observed for responders in the wild-type (Figure 4C) or Δ*comS* (Figure 4E) genetic background. These data show that, in this experimental system, ComRS signaling from the sender strain elicits a much weaker response from the responder than does the addition of synthetic XIP or was seen in the microtiter-based experiments detailed above. Since diffusion limitation cannot explain the results, we posited that the differences between this experiment and those described above could arise from a number of different factors that include a shorter experimental duration, a greater average distance between sender and responder cells in the agarose matrix than in the co-culture studies, more limited lysis of the senders (see below), or insufficient externalization of XIP by the sender under these particular conditions. Additionally, differences in XIP turnover due to the localized accumulation of a protease(s) that cleaves XIP before it can diffuse could also explain the differences between experiments.

**Figure 4.**
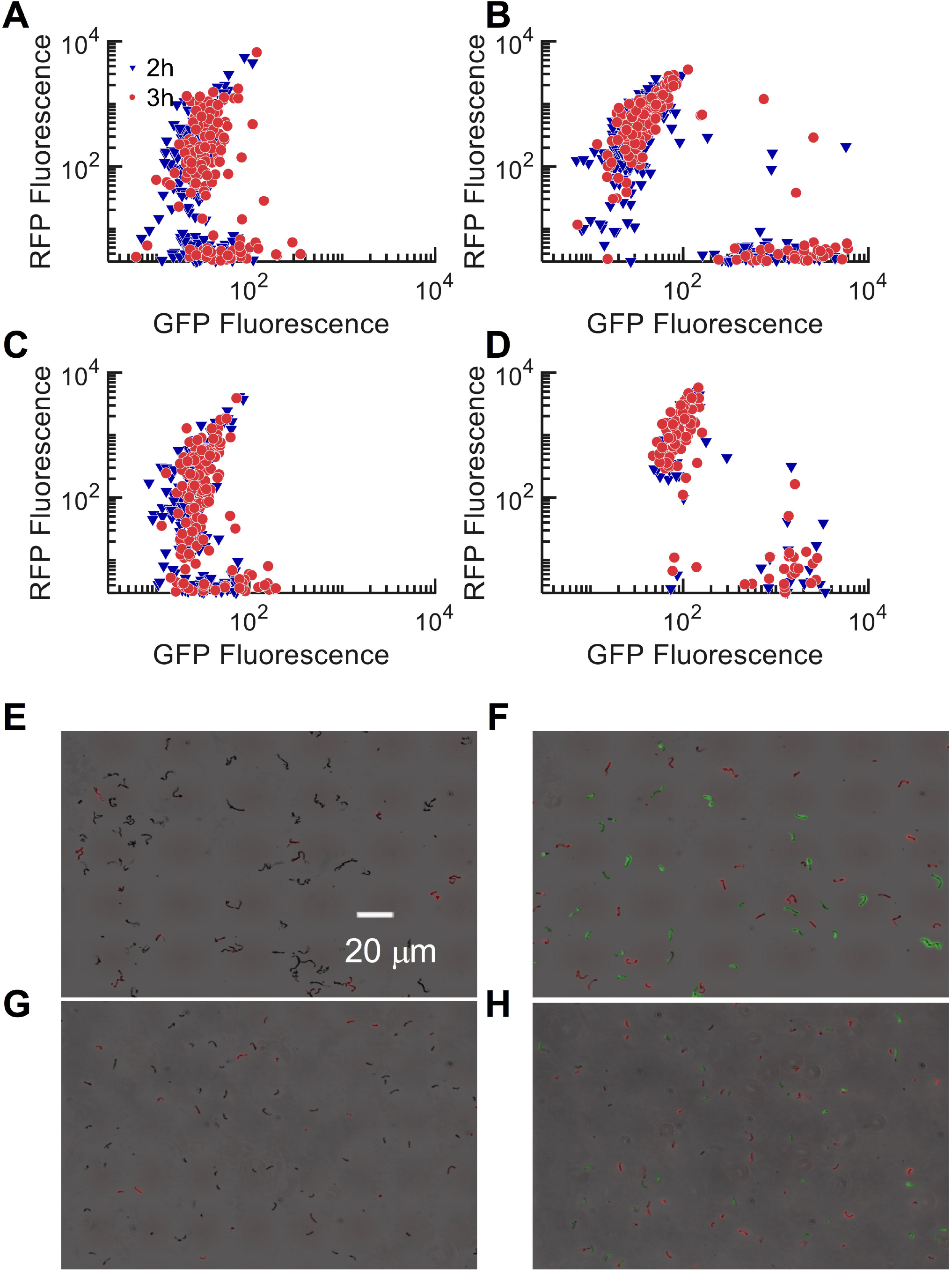
Single-cell observations of ComRS signaling in co-cultures. Co-cultures of *comS*-overexpressing sender (P*comX*:*dsRed*/pIB184*comS*) and either P*comX*::gfp/UA159 or P*comX*::gfp/ Δ*comS* responder in a low-melting-point agarose/FMC gel. After loading the gel into a microfluidic channel, cells were imaged at hourly intervals by phase contrast and fluorescence microscopy. (A) Fluorescence of P*comX*::*dsRed*/pIB184*comS* and P*comX*::gfp/UA159 co-culture at 2 h (blue; triangles) and 3 h (red; circle) time points as a single cell scatter plot. (B) P*comX*::*dsRed*/pIB184*comS* and P*comX*::gfp/UA159 co-culture at 2 h and 3 h with 1 μM sXIP injected as control. (C) P*comX*::*dsRed*/pIB184*comS* and P*comX*::gfp/Δ*comS* co-culture at 2 h and 3 h time points. (D) P*comS*::*dsRed*/pIB184*comS* and P*comX*::gfp/Δ*comS* co-culture at 2 h and 3 h time points with 1 μM sXIP injected as control. (E-H) Representative images of (A-D) in the same order.

One interesting finding from this experiment is that self-activation from the sender strain (intracellular signaling) was more apparent and of a greater magnitude than observed for the responder strain (intercellular signaling). As seen in Figure 4A and 4C, the senders are self-activating and the responders remain largely inactive, in terms of *comX* promoter activity, to the signal in comparison to when sXIP was added (Figs. 4B and 4D). Such behavior may indicate, at least within the confines of this experimental design, that the ComRS pathway is more efficient for intracellular signaling than for intercellular communication. The *S. mutans* competence pathway is unique among streptococci in that a ComCDE-like system, the activation of which is dependent on the quorum sensing molecule CSP, is linked to ComRS and that addition of sCSP to a peptide-rich medium results in a ~10^3^ increase in transformation efficiency. One must therefore consider, based on these findings, whether the ComRS pathway in *S. mutans* functions primarily as an intracellular signal driven by its positive feedback loop. Such a model for the importance of internal auto-activation by ComRS in complex medium, such as BHI, has been previously presented (9). Importantly, an ability for XIP to function in self-activation of cells and also as an intercellular signaling molecule are not mutually exclusive. There is a high degree of specificity of the *S. mutans* ComR protein for XIP from *S. mutans*, but XIP variants from other streptococcal species do not appear to interact with ComR of *S. mutans*; whereas certain other ComR proteins are able to interact with XIP from non-cognate species (21). Also of interest is the fact the ComS is extremely highly conserved between isolates of *S. mutans* (30). Perhaps the stringency of the *S. mutans* ComR-XIP interaction and high degree of conservation of XIP sequence in this organism reflects evolutionary pressures and niche adaptations that favored activation of competence only with a signal input from other strains of *S. mutans* in complex populations in humans.

### ComRS signaling in biofilm populations analyzed by microscopy and flow cytometry

As noted above, the natural environment for *S. mutans* is in biofilms, enmeshed in exopolymeric material of bacterial and host origin, so we next evaluated ComRS signaling from sender to responder in an *in vitro* biofilm model system where the exopolysaccharides were generated during the experiment by enzymatic activities of *S. mutans*. To achieve an approximately equal number of sender and responder cells in the model biofilm system for the duration of the experiments, an optimal ratio of sender:responder inoculum was first determined. Beginning with a 1:1 ratio of sender:responder grown in FMC medium and incubating the biofilms for 18 hours, it was determined after biofilm dispersal and plating that the *comS*-overexpressing strain was underrepresented with respect to the P*comX*::*gfp* responder strain, with the responder constituting 91 ± 3% of the viable colonies recovered (Supplemental Figure 2). We attributed this observation to the somewhat slower growth of the *comS* overexpressing strain (Supplemental Figure 3), and possibly to an enhancement in lytic behaviors associated with *comS* overexpression. However, since ComRS signaling is impaired in acidic conditions (28, 29), we reasoned that different inocula ratios of sender:responder coupled with replacement of the supernates with fresh medium after 6 h might enhance the representation of the *comS*-overexpressing (sender) strain in the biofilm. Allowing for sufficient sender representation was considered essential for the study of signaling. We tested three different ratios of sender:responder biofilm inocula: 4:1, 1:1, and 1: 4. The 4:1 sender:responder inoculum resulted in roughly equivalent representation of the two strains after 18 h of incubation; 48 ± 10% of the viable colonies recovered were senders. The 4:1 ratio of sender:responder also yielded the greatest amount of biomass, as measured by crystal violet staining at the end of the 18 h incubation period. Hence, all biofilm experiments were conducted with the 4:1 biofilm inoculum of sender:responder, unless otherwise noted.

ComRS signaling within co-culture biofilm populations of *S. mutans* was visualized after 18 h of incubation in FMC medium, with replacement of spent medium with fresh medium at 6 h. As a positive control, sXIP was added with the fresh medium at the 6-h time point in a final concentration of 50 nM. Fluorescence images from both the sXIP-treated control and co-culture biofilms that received no sXIP treatment were obtained for both the RFP-marked sender strain and the GFP responder strain using confocal microscopy (Figure 5A). To quantify the proportion of GFP-positive responders, biofilm populations grown under the same conditions were harvested, sonicated to isolate individual cells, and analyzed via flow cytometry. A robust response to sXIP was seen in the control biofilm population, with 65 ± 3% of single cells expressing GFP (Figure 5B). In comparison, the co-culture biofilm population had measurable GFP expression, but in a diminished proportion of the cells (12 ± 2%). In addition, the sender strain constituted a greater proportion of the co-culture population (45 ± 4%) than in biofilms treated with sXIP (18 ± 3%), likely due to enhanced sensitivity of the ComS-overproducing strain to growth inhibition and/or induction of lysis by sXIP.

**Figure 5.**
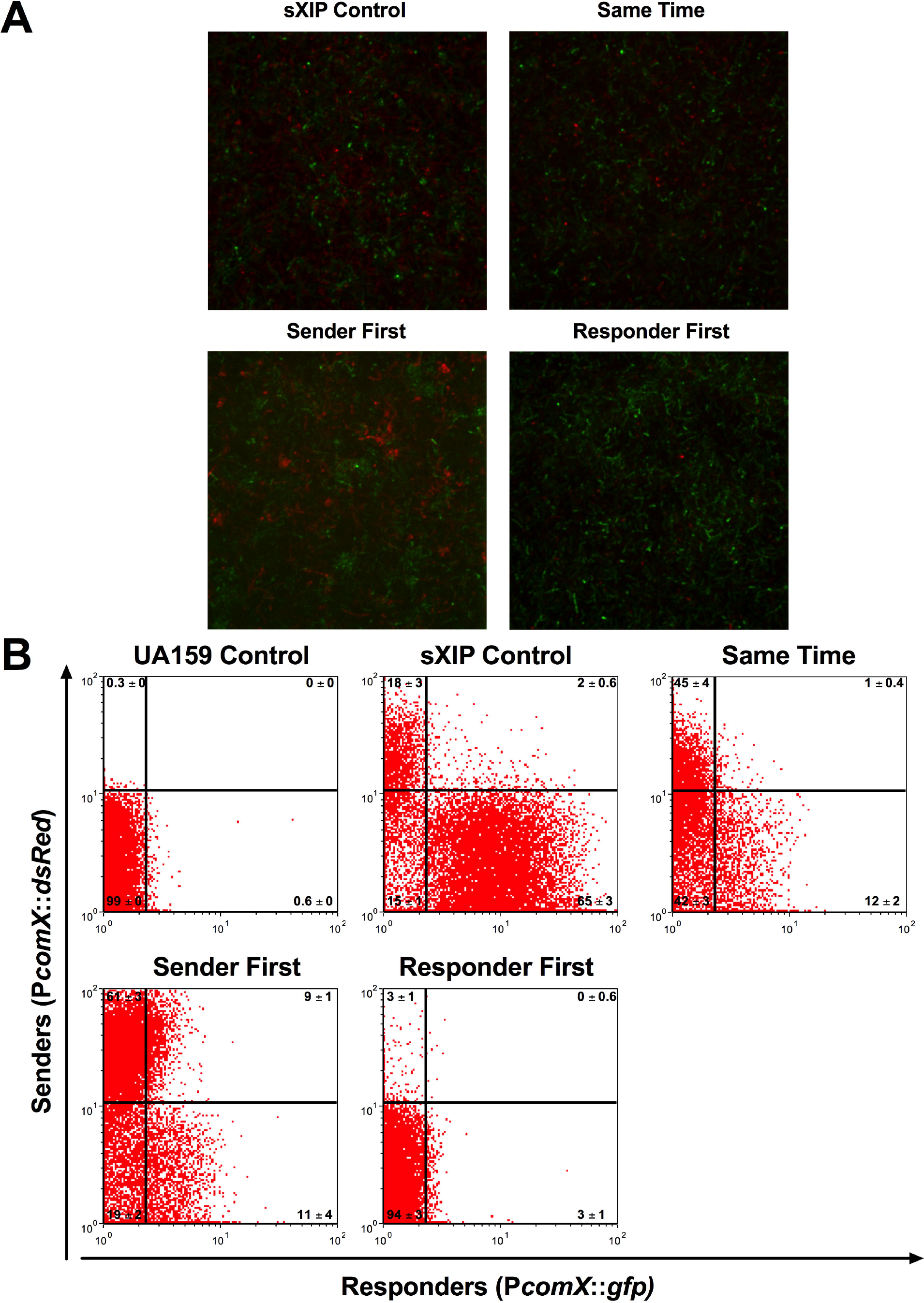
ComRS signaling in biofilms. Observation of ComRS signaling in 18 h biofilms. (A) Selected maximum intensity z-section confocal microcopy images of co-culture biofilms. Images are of a 10 µm section of fluorescent range within the biofilm, collected at 1 µm intervals using a 63X/1.40 oil objective lens. Y-axis labeling of the panel denotes the order of biofilm inoculation at time 0 h. (B) Quadrant analysis of collected flow cytometry data from similarly-grown co-culture biofilms shown in A. Y-axis shows dsRed intensity and X-axis shows GFP intensity. Quadrants are set to UA159 control. Flow cytometry data was collected from three independent experiments with triplicate samples.

As the *comS*-overexpressing (sender) strain has apparently decreased fitness in biofilm populations compared to a similar strain with an otherwise wild-type genetic background (responder), we next grew biofilms with an original inoculum of either sender or responder, and added the other strain at 6 h, along with fresh medium, to establish the co-cultures. Under growth conditions in which the *comS*-overexpressing strain was allowed to establish first in the biofilm, an increased proportion (61 ± 3%) of cells displayed *dsRed* fluorescence by flow cytometry, compared to when the strains were added together in the initial inoculum (45 ± 4%). However, the proportion of the GFP-positive PcomX::*gfp* strain remained unchanged when contrasting co-inoculation with inoculation with the responder strain at 6 h (11 ± 4% to 12 ± 2%, respectively). However, when the responder strain was established first, very little GFP or RFP fluorescence activity was observed, either by confocal microscopy or flow cytometry. This is most likely due to the *comS*-overexpressing strain not being able to establish and/or persist after the strain with the wild-type genetic background had already established a biofilm. Overall, ComRS signaling in co-culture biofilms was strongly dependent on the timing of introduction of the second strain and the proportions of the sender; a finding that is not surprising in light of the established impacts of low pH, growth phase and other factors on the efficiency of XIP-dependent activation of *comX.*

### Sucrose reduces intercellular XIP signaling in biofilms

*S. mutans* strains are genomically and phenotypically diverse members of the oral microbiome that predominantly colonize hard surfaces. A substantial body of evidence implicates these organisms as primary etiological agents of human dental caries (31). Among the many attributes that enable *S. mutans* to be an effective caries pathogen are its potent acidogenic and aciduric properties, coupled with its capacity to utilize sucrose to form copious quantities of extracellular polysaccharide (EPS) via three glucosyl- and one fructosyl-transferase enzymes (Gtfs and Ftfs) (32). Gtfs, through *in situ* synthesis on the acquired enamel pellicle, provide initial colonization sites for *S. mutans*, and produce the insoluble EPS matrix that encases the bacteria, leading to the establishment of complex 3-dimensional biofilm structures (33). Within these structures are highly diverse microenvironments that may influence diffusion of chemical compounds and peptides that function in interbacterial communication. All experiments herein were, to this point, conducted using FMC containing 20 mM glucose as the sole carbohydrate source. To explore how addition of sucrose and production of an EPS matrix would impact XIP signaling, we grew co-culture biofilms with low (2.5 mM to 15 mM), medium (5mM to 10 mM) and high (9 mM to 2 mM) ratios of sucrose to glucose; sucrose is a dissacharide, so the weight-to-volume concentrations of fermentable carbohydrates were constant across all experiments. In all conditions in which sucrose was present, XIP activation within the co-culture biofilm was evident (Figure 6). Interestingly, though, when sucrose was provided, a spatial organization pattern between the *comS*-overexpressing sender and PcomX::*gfp*/UA159 responder became apparent, with the two strains segregating into different regions of the biofilms (Figure 6A). In terms of both proportion of cells responding and overall *gfp* intensity from the PcomX::*gfp* strain, measureable *gfp* fluorescence was reduced under conditions in which sucrose was added to the growth medium (Figures 6B). In 20 mM glucose alone, the proportion of *gfp* expressers was 19 ± 2% and median GFP intensity was 4.3 ± 0.5 au (arbitrary fluorescent units), whereas 5 ± 1% of the population expressed *gfp* and intensity was recorded as 3.1 ± 0.1 au under all sucrose conditions tested. Similar to the co-culture experiments within the FMC/agarose gel, an increase in the strength of intracellular signaling was observed as RFP intensity measured in the sender strain increased from 17.9 ± 0.9 au in biofilms formed in 20 mM glucose to 25.3 ± 0.9 au when the highest sucrose concentration was present. Similar decreases in measureable *gfp* fluorescence due to the presence of sucrose were observed when the co-culture experiment was conducted in planktonic growth conditions rather than biofilms (Figure 6C). The abolishment of XIP signaling in medium- and high-sucrose conditions was not due to changes in the *comS*-producing sender cells, as P*comS* feedback showed similar activation in all conditions (Figure 6D). Together the biofilm and planktonic experiments show that addition of sucrose to the growth medium significantly impact the ability of the sender strain to activate the responder.

**Figure 6.**
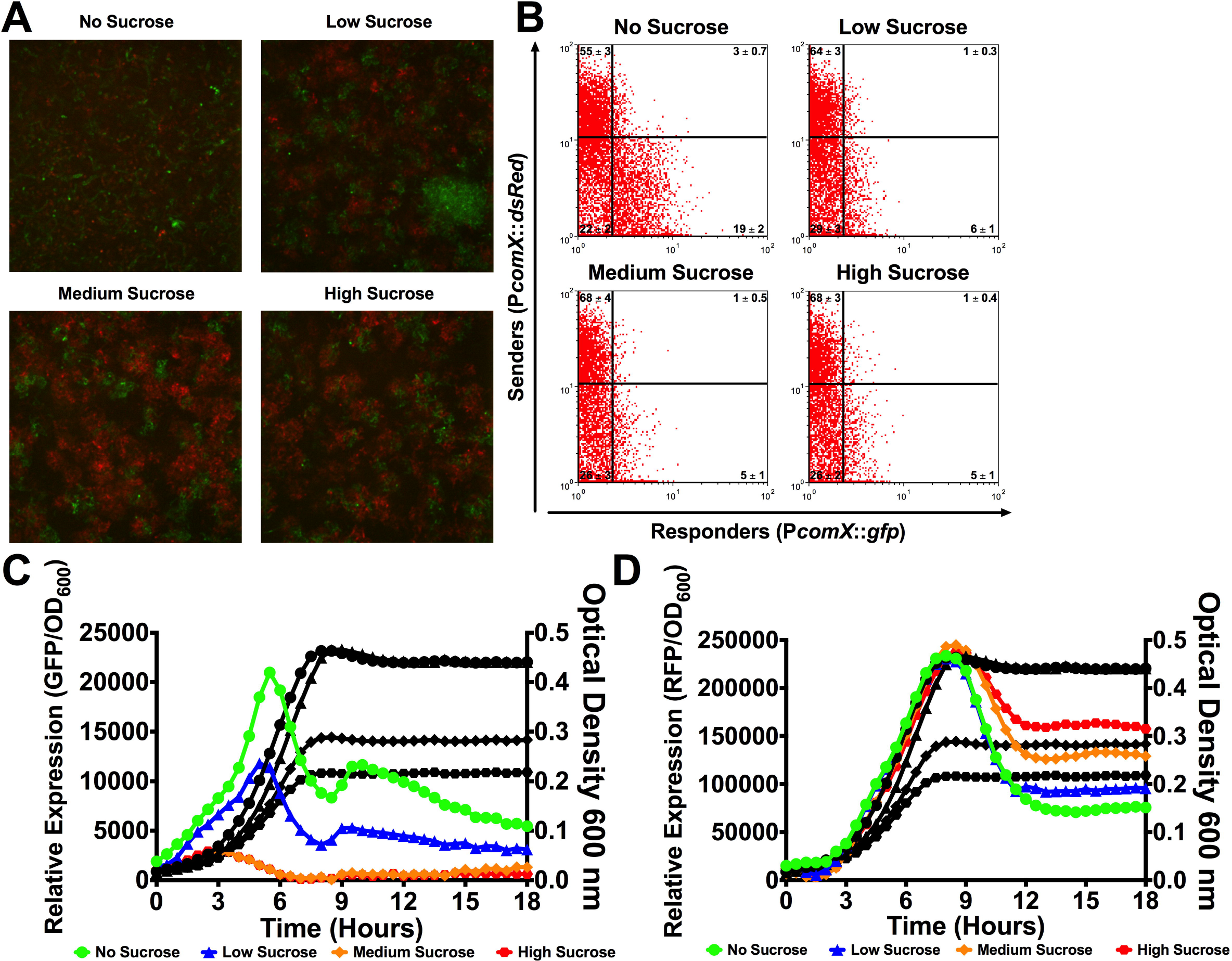
Impact of sucrose on ComRS signaling in biofilms. Observation of ComRS signaling in 18-h biofilms grown in different concentrations of sucrose. (A) Selected maximum intensity z-section confocal microcopy images of co-culture biofilms. Images are a 10 µm section of fluorescent range within the biofilm, collected at 1 µm intervals using an 63X/1.40 oil objective lens. X-axis labeling of the panel denotes amount of carbohydrate source used in biofilm growth medium (No sucrose = 20 mM glucose; low sucrose = 15 mM glucose and 2.5 mM sucrose; medium sucrose = 10 mM glucose and 5 mM sucrose; high sucrose = 2 mM glucose and 9 mM sucrose). As sucrose is a disaccharide, carbohydrate concentration (w/v) was the same in each condition. (B) Quadrant analysis of collected flow cytometry data from similarly-grown co-culture biofilms as shown in A. Y-axis shows dsRed intensity and X-axis shows GFP intensity. Quadrants are set to UA159 control. Flow cytometry data was collected from three independent experiments with triplicate samples. (C) Relative GFP expression and (D) relative RFP expression with OD_600_ measurements during co-culture growth of pIB184comS/UA159 and P*comX*::*gfp*/UA159 either no sucrose (green; circles), low sucrose (blue; triangles), medium sucrose (orange; diamonds), or high sucrose (red; squares) added to the growth medium as a carbohydrate source. Each assay was performed with biological triplicates.

To further evaluate the basis for the negative impact of sucrose and the resultant polysaccharide matrix on intercellular signaling by XIP within biofilm populations, biofilms of P*comX*::*gfp*/UA159 were grown with low, medium, and high ratios of sucrose to glucose for 5 h before sXIP was added. GFP production was monitored at selected time intervals, both by relative fluorescence in a plate reader and by confocal microscopy (Supplemental Figure 4). Ample *comX* activation was noted in biofilms cultured in low-sucrose conditions when a final concentration of 200 nM or 2 µM sXIP was provided. However, at the intermediate sucrose concentration, activation was only seen with 2 µM sXIP and yielded a relative fluorescence that was 3.5-fold lower compared to the low-sucrose condition. The most substantial effects were seen in the high-sucrose condition, where 2 µM sXIP was unable to activate the P*comX* responder strain. In fact, 10 µM sXIP was needed to measure GFP production within the high-sucrose biofilms. Collectively, these data highlight that provision of sucrose, which dramatically alters the biofilm EPS matrix and influences the physiology and transformability of *S. mutans* (34, 35), has an overall negative impact on the response of cells to XIP.

One potential explanation for the reduced response by the responder strain when various amounts of sucrose were present in the growth medium is slower diffusion of the secreted XIP peptide within the biofilms due to increased EPS formation. The reduced diffusion of XIP could lead to changes in the spatial distribution and temporal dynamics of ComRS signaling within biofilm populations, leading to increased phenotypic heterogeneity within the biofilm. Such observations were recently noted in the study of quorum sensing systems within biofilms under flowing conditions (36). The images in Figure 6A show that some spatial correlation between sender and responder is evident, such that clusters of senders and responders appear segregated, even though the biofilm inoculum consists of uniform suspension of senders and responders. A comparison of merged and brightfield images also shows that certain cells within the biofilm population remain unresponsive to the sender’s signal (Supplemental Figure 5). Similar phenotypic heterogeneity was recently noted in monoculture biofilms that had been treated with sXIP peptide, and notably, there was little evidence of cell death in biofilms cultured under similar conditions (24). Thus, it will be interesting to unravel why these cells within the biofilm are unresponsive, although it can be hypothesized that some are unresponsive due to slow growth or to being confined to microenvironments that inhibit uptake or responses to signal(s) inputs. Further study of the spatial organization of various modified sender and responder strains should yield a more complete understanding the role of biofilm architecture on signaling, and vice versa.

### Lysis of the sender strain contributes significantly to XIP signaling

A critical gap in our current understanding of the ComS/XIP system of *S. mutans* is that the export apparatus and protease that generate released XIP from ComS have not been identified. Several attempts have been made to identify both the presumptive exporter and the protease in *S. mutans* (13, 20, 37). The ABC transporter PptAB was identified recently in *S. pyogenes* as an SHP exporter, but XIP secretion in *S. mutans* was only partially reduced in a *pptAB* deletion (38). These results indicate that other mechanisms exist by which XIP appears in the supernatant fraction of *S. mutans* cultures (13, 15). Since the studies detailed above provide ample evidence for intercellular activation of ComRS under conditions in which substantial amounts of cell lysis occur, such as overnight planktonic growth (Figure 2) and growth within biofilms (Figures 5 and 6), but a lack of intercellular activation when early exponential phase cells were spatially separated and fixed within an agarose suspension (Figure 4), we examined whether active cell lysis was important for the release of the ComS/XIP signal to potential responders. To accomplish this, we monitored XIP signaling activity in supernates of *comS*-overexpressing cells in a wild-type genetic background and in a strain lacking the major autolysin, AtlA (Δ *atlA*) (Figure 7) (39). Whereas supernates from UA159 or the Δ *atlA* mutant failed to activate the P*comX*::*gfp*/UA159 reporter, the supernates from the *comS*-overexpressing strain in an AtlA-positive strain activated P*comX* expression in the responder, similar to the results reported above (Figure 2A). However, no detectable GFP fluorescence was observed when the supernates from the *comS*-overexpressing strain carrying the *atlA* deletion was supplied to the responder strain (Figure 7A). These data are corroborated by growth curves of the P*comX*::*gfp*/UA159 responder strain in the selected supernates, as growth in the supernates from the 184*comS*/UA159 strain was slower than in supernates from UA159, Δ*atlA*, attributable to diminished XIP-dependent growth inhibition and/or XIP-induced lysis of the responder. Experiments using a P*comX*::*gfp*/Δ which eliminates the auto-feedback loop and self-activation through activation of *comS* by ComR-XIP, also showed no activation in response to the *atlA* mutant supernates (Figures 7B). These data provide intriguing new evidence that externalization of ComS/XIP in *S. mutans* may depend, entirely or in part, on cell lysis or loss of membrane integrity, rather than on a dedicated secretion apparatus, as is the case for a number of other streptococci (16, 38).

**Figure 7.**
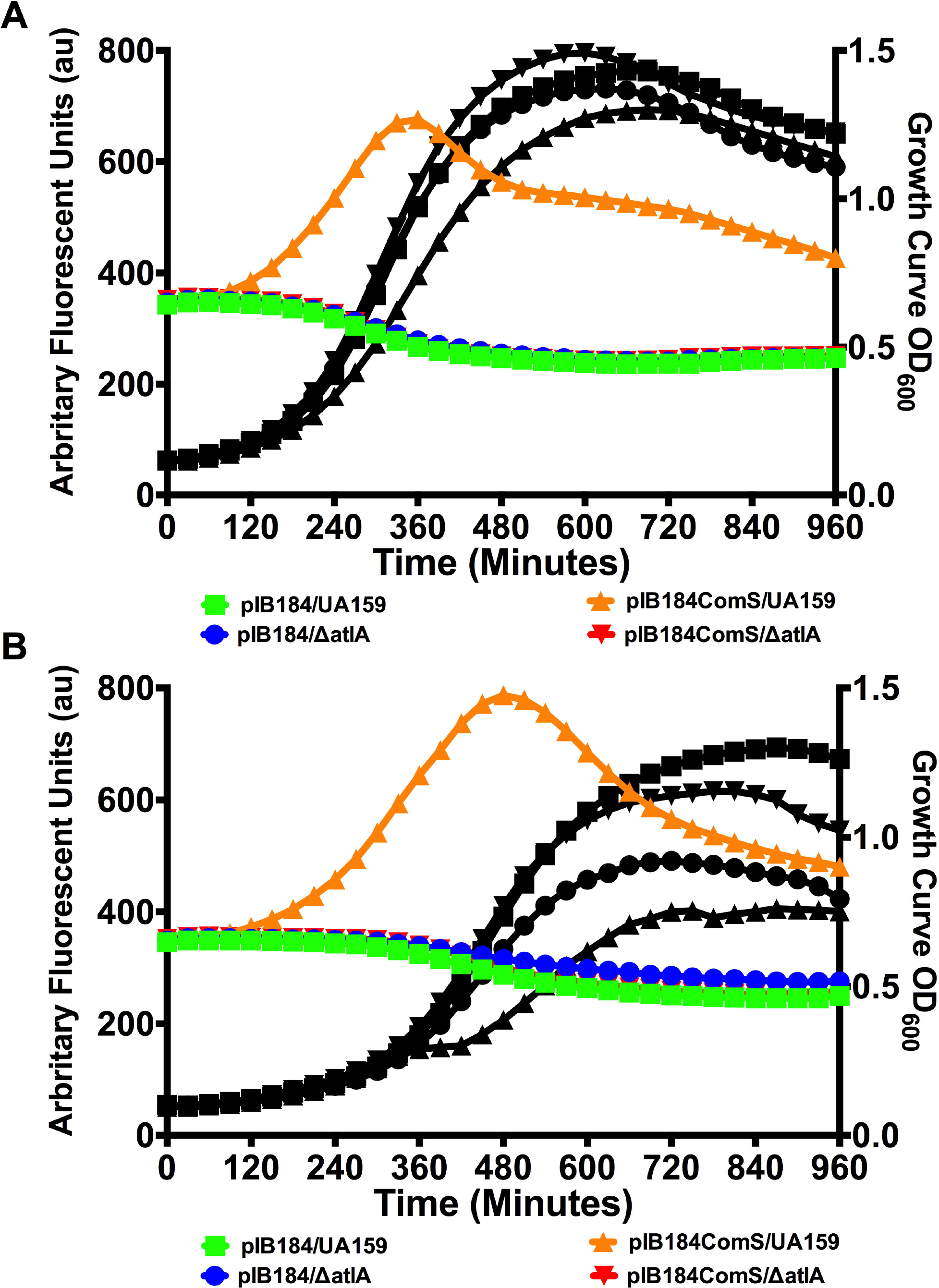
Impact of cell lysis on ComRS Signaling. Fluorescence reporter activity in arbitrary fluorescent units (au; colored lines) and growth curves (black lines) of responder strains grown in supernates of sender strains. (A) GFP fluorescence reported as arbitrary fluorescent units of P*comX*::*gfp*/UA159 reporter cells grown in overnight supernates (filtrates) of either pIB184/UA159 (green; squares), pIB184/Δ*atlA* (blue; circles), pIB184*comS*/UA159 (orange; upward triangles), or pIB184*comS*/Δ*atlA* (red; downward triangles). Overnight supernates were pH-corrected to 7.0 using 6.25N NaOH and glucose was replenished to a concentration equivalent to an additional 20 mM before filtering the supernatant fluids through a 0.22 μM PVDF syringe. (B) GFP fluorescence of P*comX*::*gfp*/Δ*comS* reporter cells grown in overnight supernates (filtrates) as in A. Each assay was performed with biological triplicates.

A number of properties of the ComRS system of *S. mutans* in particular may provide an explanation for why cell lysis could be the primary pathway for XIP externalization. First, ComRS is extremely highly conserved among *S. mutans* isolates, with almost no variation in ComS sequence and none in the sequence of XIP (30). Similarly, ComR of *S. mutans* very specifically recognizes *S. mutans* XIP, but not XIP from other sources. Similar to the ComS sequences of the *Streptococcus bovis* group (40), *S. mutans* ComS sequence is short (17-aa) and lacks a typical signal sequence used for active export. It is also notable that *S. mutans* has an obligatory biofilm lifestyle, where regulated cell death and lysis, along with eDNA release (41), are critically important for biofilm maturation and stability (42). *S. mutans* may thus have evolved in a way that a dedicated ComS/XIP exporter was dispensable, as autolysis of bacterial cells during biofilm growth could be sufficient to release XIP at levels that are effective for its function(s) (41). The release by lysis also would serve as an altruistic signal to viable *S. mutans* that *S. mutans* DNA has been released into the environment, which may have aided in the diversification of the species by tuning competence to the preferential assimilation of DNA that could be sufficiently homologous to recombine with the competent population. If confirmed, these findings may also explain why *S. mutans* bacteriocin production and autolysis are more intimately intertwined with competence development than in some other streptococcal species (43, 44). CSP-induced activation of bacteriocins, some of which can kill sensitive strains of *S. mutans*, could trigger lysis in the sensitive sub-population, which would then release XIP and DNA for the now-competent members of the biofilm. The hypothesis also takes into consideration Wenderska *et. al.*’s observation that deletion of *comX* reduces the abundance of XIP in culture supernates (37), as this deletion would remove competence-driven lysis as a release pathway. It is also interesting to posit that maturation of ComS to XIP may actually be a consequence of induced cell lysis and activation of proteolytic pathways, such that when a cell lyses, XIP would be released. Currently unknown protease(s), which may be activated during programmed cell lysis, could then generate the 7-aa XIP peptide from the 17-aa ComS during cell death to activate nearby responder cells, allowing for the uptake of homologous pieces of DNA leading to enhanced biofilm formation and an increased fitness of the present population.

## SUMMARY

Recent analysis of bacterial communication at the single cell level and *in situ* in biofilm systems has led to revision of numerous views of intercellular signaling in “natural” populations. While many of these advances have been made with well-established model organisms (e.g. *Bacillus subtilis, Staphylococcus aureus, Escherichia coli, Pseudomonas aeruginosa*), the oral pathogen *S. mutans* regulates multiple biological processes, including genetic competence, bacteriocin production, biofilm maturation and tolerance of environmental insults, through a suite of small hydrophobic peptides (26, 27) and dedicated transcriptional regulators (45, 46). Many of these are unique to this organism, function differently than in paradigm organisms, and are critical to the ability of *S. mutans* to cause disease.

Overall, the work presented here highlights a new model system for studying Gram-positive bacterial communication based on naturally-derived signal oligopeptides from an overexpressing population, as opposed to the addition of synthetic peptide(s). The model shows that XIP can, in fact, act as an intercellular signal that does not require cell-cell contact. The model also allows investigation of responses to XIP in a biological setting that more closely mimics the natural growth environment of the organism. This work also emphasizes the differences in signaling behaviors that occur between planktonically grown cultures and those that are grown within biofilms, where spatial and temporal distribution patterns can have a more significant impact on interpretation of signal inputs. Development of this system facilitates the modelling of ComRS behaviors within an environment such as dental plaque, where competition of *S. mutans* with commensal streptococci strongly influences the pathogenic potential of the biofilms (47). Importantly, this work also highlights for the first time the connection between lysis and XIP release, offering an alternative hypothesis for why XIP has been detected mainly in culture supernates. Future investigations can provide insight into how signaling impacts genetic and physiological control of early biofilm growth, responses to environmental stresses and competition between competing oral species, ultimately yielding information that can be used to disrupt key components of the signaling circuit to decrease the proportions of pathogens in dental biofilms.

## EXPERIMENTAL PROCEDURES

### Bacterial strains and growth conditions

*S. mutans* wild-type strain UA159 and its derivatives (Table 1) were grown in either brain heart infusion (BHI) (Difco) or FMC medium (48) that was supplemented with 10 µg ml^−1^ erythromycin and 1 mg ml^−1^ of kanamycin or spectinomycin, as needed. Unless otherwise noted, cultures were grown overnight in BHI medium with the indicated antibiotics, if needed, at 37°C in a 5% CO_2_, aerobic atmosphere. The next day, cultures were harvested by centrifugation, washed twice in 1 mL of phosphate-buffered saline (PBS), and resuspended in PBS to remove all traces of BHI. PBS cell suspensions were then transferred to 5 mL polystyrene round-bottom tubes (Corning Incorporated). The samples were then sonicated using a Fisher Scientific Model 120 Sonic Dismembrator in the water bath mode at 100% amplitude for three intervals of 30 s each, with placement on ice for the intervals. Sonicated cell suspensions were subjected to a final centrifugation to remove any cellular debris and resuspended in the desired medium before diluting to begin each experiment. Synthetic XIP (sXIP, aa sequence = GLDWWSL), corresponding to residues 11-17 of ComS, was synthesized and purified to 96% homogeneity by NeoBioSci (Cambridge, MA). The lyophilized sXIP was reconstituted with 99.7% dimethyl sulfoxide (DMSO) to a final concentration of 2 mM and stored in 100 μL aliquots at −20°C

**TABLE 1.** List of strains

| Strain or plasmid | Relevant Characteristics* | Source or Reference |
| --- | --- | --- |
| *S. mutans* Strains |  |  |
| UA159 | Wild-type | ATCC 700610 |
| *PcomX::gfp*/ UA159 | UA159 harboring *PcomX::gfp* | (9) |
| *PcomX::gfp/ΔcomS* | *ΔcomS* harboring *PcomX::gfp* | (9) |
| *PcomX::gfp/ΔoppA* | *ΔoppA* harboring *PcomX::gfp* | (9) |
| *PcomX::gfp*/pIB184 | UA159 harboring *PcomX::gfp* and pIB184 | (28) |
| *PcomX::gfp*/pPMZ | UA159 harboring *PcomX::gfp* and pPMZ | This study |
| pIB184comS/ UA159 | UA159 harboring pIB184comS | (28) |
| *PcomS::dsRed*/pIB184comS | UA159 harboring *PcomS::dsRed* and pIB184comS | This study |
| *PcomX::dsRed*/pIB184comS | UA159 harboring *PcomX::dsRed* and pIB184comS | This study |
| *ΔatlA* | *atlA* (SMU.704c) :: NPKm<sup>R</sup> | (39) |
| pIB184comS / *ΔatlA* | *ΔatlA* harboring pIB184comS | This study |
| Plasmids |  |  |
| pDL278 | *E. coli* - *Streptococcus* shuttle vector, Sp<sup>R</sup> | (51) |
| pIB184 | Shuttle expression plasmid with the constitutive P23 promoter, Em<sup>R</sup> | (25) |
| pPMZ | LacZ fusion integration vector based on pMC195 and pMC340B; Km<sup>R</sup> | (52) |

\* Em, erythromycin; Km, kanamycin; Sp, spectinomycin.

### Construction of bacterial strains

Mutant strains of *S. mutans* were created using a PCR ligation mutagenesis approach, as previously described (49). Overexpression of genes was achieved by amplifying the structural genes of interest from *S. mutans* UA159 and cloning into the expression plasmid pIB184 (25). Transformants were confirmed by PCR and sequencing after selection on BHI agar with appropriate antibiotics. Plasmid DNA was isolated from *E. coli* using QIAGEN (Chatsworth, Calif.) columns, and restriction and DNA-modifying enzymes were obtained from Invitrogen (Gaithersburg, Md.) or New England Biolabs (Beverly, Mass.). PCRs were carried out with 100 ng of chromosomal DNA by using *Taq* DNA polymerase, and PCR products were purified with the QIAquick kit (QIAGEN).

### Measurements of GFP fluorescence with plate reader

For measurements of GFP fluorescence, co-cultures were inoculated from washed and sonicated overnight cultures in FMC medium at a 1:1 ratio, unless noted otherwise. Inoculated medium (175 μL) was added to each well along with a 50 μL mineral oil overlay in a Costar^™^ 96 well assay plate (black plate with clear bottom; Corning Incorporated) and incubated at 37°C. At intervals of 30 minutes for a total of 18 hours, absorbance at 600 nm along with GFP fluorescence (excitation 485/20 nm, emission 528/20 nm) was measured with a Synergy 2 multimode microplate reader (BioTek). Relative expression was calculated by subtracting the background fluorescence of UA159 (mean from six replicates) from raw fluorescence units of the reporter strains and then dividing by OD_600._

### Measurement of diffusive XIP spreading

To estimate a diffusion constant for sXIP and the ComRS system, P*comX*::*gf*p/ Δ *comS* cells were injected into an IBIDI microslide (ibidi GmbH, µ-slide VI), a slide having 6 parallel, narrow channels, each with a loading port at both ends. The cells were allowed to settle to the surface of the channels before a 2% low-melting-point agarose/FMC gel was pushed through each channel to immobilize individual cells on the channel window. Injection of the agarose/FMC mixture removed any cells that were not stuck to the slide. sXIP that was reconstituted in DMSO and diluted into FMC to a final 1 µM concentration that was then deposited in one port of each channel, with an equal volume of DI water in the opposite port to balance the hydrostatic pressure in the channel. The resulting culture was then incubated at 37°C with mineral oil sealing the loading ports to minimize drying of the cooled agarose. Green fluorescence of individual P*comX*::*gf*p/ Δ *comS* reporter cells was measured as a function of position and time, relative to the XIP loading, in order to estimate the diffusion constant for sXIP in the medium. Cells were imaged at 20X (CFI Plan Fluor DLL, NA 0.5, Nikon) onto a cooled CCD camera (CoolSNAP HQ2, Photometrics). Images were collected in phase contrast, GFP fluorescence (using a Nikon C-FL GFP HC HISN zero shift filter cube), or red fluorescence (C-FL Y-2E/C Texas Red filter cube, Nikon). Three images of each channel were collected at intervals of one hour, for periods up to five hours, after introduction of sXIP. The expression of GFP in individual cells was quantified by a previously described method (9, 50). Fitting of the diffusion coefficient was performed in Matlab^®^ (The Mathworks, Inc.) and was done by minimizing the difference between measured GFP response and one calculated from a 1D diffusion equation model with a delta function of XIP concentration at the origin as an initial condition. The error on this fit was checked by a 1000-iteration bootstrap method, with the standard deviation of calculated diffusion coefficient values used as the fit error.

### Co-culture within agarose/FMC mixture

P*comX*::*dsRed*/pIB184*comS* (sender strain) and P*comX*::*gfp* reporter in either a UA159 or Δ *comS* background (responder strain) were mixed together at OD_600_ of 0.1 in liquid FMC at a 2.3::1 ratio before injection into an IBIDI microslide and immobilized with a 2% low-melting temperature agarose/FMC mixture. The resulting culture was grown in an incubated chamber with mineral oil on top of the injection holes to minimize drying. Imaging was performed at intervals to examine reporter fluorescence, as previously described (9, 29). sXIP (1 µM) was added as a positive control, and channels containing solely GFP reporter cells were also imaged as a negative control.

### Fitness assessment

Fitness between the *comS*-overexpressing strain and the PcomX::*gfp* responder strain carrying a kanamycin resistance marker (pPMZ) was determined from 18 h biofilms inoculated with either a 4::1, 1::1, or 1::4 ratio of sender::responder and grown in 6 well polystyrene flat bottom plates (Corning Incorporated). At the end of the 18 h incubation, biofilms were washed twice in PBS to remove unattached cells, scraped from the 6 well plates using a cell scraper, and sonicated in a water bath sonicator for 3 intervals of 30 seconds in 5 mL polystyrene round-bottom tubes. Single cells resuspened in PBS were then serially diluted and plated on either BHI, erythromycin-BHI, or kanamycin-BHI plates for selective plating. After 48 hours of incubation at 37°C in 5% CO_2_, plates were removed, imaged, and viable colonies enumerated from each plate. Fitness of each strain was then determined by taking the sum of viable colonies from the erythromycin-BHI and kanamycin-BHI plates, dividing by the erythromycin-BHI count and multiplying by 100 to determine a viable count percentage returned of each strain from the grown biofilm.

### Confocal laser scanning microscopy

Sonicated overnight cultures were washed and re-suspended in FMC before being diluted 1:50 overall into fresh medium with the desired sender:responder co-culture ratio. Diluted cell suspensions (350 µL) were inoculated into each well of an 8-well µ-Slide (ibidi USA) chambered coverslip as well as a 6-well polystyrene flat bottom plate (2.5 mL) to be used for flow cytometry analysis. Plates were incubated at 37°C in a 5% CO_2_, aerobic atmosphere for a total of 18 h. At the 6-h time point, spent medium was removed from the biofilms and fresh medium applied along with either sXIP or another strain, if desired. Prior to analysis by microscopy, wells were washed 3 times with PBS and were kept hydrated with 100 µL of PBS. Biofilm images were acquired using a spinning disk confocal system connected to a Leica DMIRB inverted fluorescence microscope equipped with a Photometrics cascade-cooled EMCCD camera. GFP fluorescence was detected by excitation at 488 nm and emission was collected using a 525 nm (±25 nm) bandpass filter. Detection of dsRed fluorescence (RFP) was performed using a 642-nm excitation laser and a 695-nm (±53-nm) bandpass filter. All z-sections were collected at 1 µm intervals using an 63X/1.40 oil objective lens. Image acquisition and processing was performed using VoxCell (VisiTech International).

### Flow cytometry

Biofilms grown in 6-well flat bottom plates were scraped from the surface of wells before being run through a FACSCalibur™ (BD Biosciences) flow cytometer. At the end of the 18-h incubation, biofilms were washed twice in PBS to remove unattached cells, scraped from the 6-well plates using a cell scraper, and sonicated in a water bath sonicator for 3 intervals of 30 seconds in 5 mL polystyrene round-bottom tubes to achieve primarily single cells for analysis. Forward and side scatter signals were set stringently to allow sorting of single cells. In total, 5 x 10^4^ cells were counted from each event, at a maximum rate of 2 x 10^3^ cells per second, and each experiment was performed in triplicate. Detection of GFP fluorescence was through a 530 nm (± 30 nm) bandpass filter, and dsRed was detected using a 670-nm long pass filter. Data were acquired for unstained cells and single-color positive controls so that data collection parameters could be properly set. The data were collected using Cell Quest Pro (BD Biosciences) and analyzed with FCS Express 4 (De Novo Software). Gating for quadrant analysis was selected by using a dot density plot with forward and side scatter, with gates set to capture the densest section of the plot. Graphing and statistical analyses were performed using Prism (GraphPad Software). *x*- and *y*-axis data represent logarithmic scales of fluorescent intensity (arbitrary units).

### Monitoring XIP-signaling using filtered supernates

Filtered culture supernates from selected strains of *S. mutans* were produced by centrifugation of cultures diluted 20-fold from overnight and grown to OD = 0.85-1.0. The resulting supernatant fluid was adjusted to pH 7.0 using 6.25 N NaOH and concentrated glucose was added to increase the amount of glucose in the supernates by 20 mM. The resulting solution was then filtered using a 0.22 μm PVDF syringe filter. P*comX*::*gfp* reporter cells with either UA159 or Δ *comS* background were diluted 20-fold from an overnight culture, grown to OD_600_ of 0.1 and collected via centrifugation. The supernates of the reporter cells were removed and replaced with the same volume of the pH- and glucose-corrected supernates of the strains of interest. For sXIP controls, supernates were replaced with fresh FMC, after which sXIP was added in the indicated concentrations. Two mL of the resulting reporter cell-supernate combinations were then placed in a Falcon 24 well plate (Corning Inc.), covered with mineral oil and the OD_600_ and green fluorescence (excitation 485nm, emission 528nm filter set) were measured in a BioTek Synergy 2^^®^^ plate reader, shaking gently before each reading to prevent cell settling.

## ACKNOWLEDGEMENTS

Research reported in this publication was supported by the National Institute of Dental and Craniofacial Research of the National Institutes of Health under Award Numbers R01 DE023339, R01 DE13239, T90 DE21990 and F31 DE024416.

